# Identification of Dynamic Microbial Signatures in Longitudinal Studies

**DOI:** 10.1101/2022.04.25.489415

**Authors:** M.Luz Calle, Antoni Susin

## Abstract

The study of microbiome dynamics is key for unveiling the role of the microbiome in human health. Addressing the compositional structure of microbiome data is particularly critical in longitudinal studies where compositions measured at different times can yield to different subcompositions.

We propose a new compositional data analysis (CoDA) algorithm for inferring dynamic microbial signatures. The algorithm performs penalized regression over the summary of the log-ratio trajectories (the area under these trajectories) and the inferred microbial signature is expressed as a log-contrast model. Graphical representations of the results are provided to facilitate the interpretation of the analysis: plot of the log-ratio trajectories, plot of the signature and plot of the prediction accuracy of the model. The new proposal is illustrated with data on the developing microbiome of infants.

The algorithm is implemented in the R package “code4microbiome” (https://cran.r-project.org/web/packages/coda4microbiome/) that is accompanied with a vignette with a detailed description of the functions. The website of the project contains several tutorials: https://malucalle.github.io/coda4microbiome/

## 1. Introduction

Microbiome composition is dynamic and the study of microbiome changes over time is of primary importance for understanding the relationship between microbiome and human phenotypes. Longitudinal studies are costly, both economically and logistically, but there is growing evidence of the limitations of cross-sectional studies for providing a full picture of the role of the microbime in human health. Microbiome longitudinal studies can be very valuable in this context, provided appropriate methods of analysis are used (Schmidt et al. 2018)

Microbiome data analysis is challenging because, among other things, the compositional nature of the data (Susin et al. 2020, Calle 2019, Gloor et al. 2016, 2017, Gloor and Reid, 2016). This is particularly critical in the context of microbiome longitudinal studies where compositions measured at different times can be affected by distinct batch effects and similar quality control or filtering protocols may yield to different subcompositions at each time point.

The log-ratio approach (Aitchison 1986), that consists in analyzing the abundances of some taxa relative to the abundances of other taxa, is subcompositionally coherent and provides an especially interesting standpoint for exploring microbiome dynamics. In longitudinal studies, the log-ratio between two groups of taxa measured at different time points gives a curve profile or trajectory for each sample. We propose to explore the association between the phenotype of interest and the shape of the log-ratio microbiome trajectories.

Among the questions outstanding about microbiome dynamics, we focus on inferring dynamic microbial signatures and propose a novel algorithm to identify a set of microbial taxa whose joint dynamics is associated with the phenotype of interest. For binary outcomes, such as disease status, we aim to identify two groups of taxa with clearly different log-ratio trajectories for cases and controls.

The algorithm performs variable selection through penalized regression over the summary of the log-ratio trajectories (the area under these trajectories). The inferred microbial signature is expressed as a log-contrast model (Aitchison, J. and Bacon-Shone,J. 1984), i.e. a log-linear model with the constraint that the sum of the coefficients is equal to zero. The zero-sum constraint ensures the invariance principle required for compositional data analysis.

The interpretability of results is of major importance in the context of microbiome studies. We provide several graphical representations of the results that facilitate the interpretation of the analysis: plot of the log-ratio trajectories, plot of the signature (selected taxa and coefficients) and plot of the prediction accuracy of the model.

The methodology is illustrated with data from the “Early childhood and the microbiome (ECAM) study” (Bokulich et al. 2016).

## 2. Materials and methods

We first describe the analysis of log-ratios between two taxa A and B in longitudinal studies, which involves the summary of the log-ratio trajectories. Then we explain how to generalize the analysis of pairwise log-ratios to identify microbial signatures involving more than two taxa.

### Log-ratio analysis and taxa prioritization

Assume *n* subjects with fixed phenotype *Y* = (*Y*_1_,…,*Y_n_*). Subject *i* has been observed in *L_i_* time points, (*t*_*i*1_, *t*_*i*2_,…, *t*_*iL*_i__). We denote by *X_i_*(*t_ij_*) = (*X*_*i*1_(*t_ij_*), *X*_*i*2_(*t_ij_*),…, *X_ik_*(*t_ij_*)) the microbiome composition of subject *i* at time *t_ij_*, where *K* is the number of taxa which is assumed to be the same for all the individuals and all the time points. *X_i_*(*t_ij_*) can represent either relative abundances (proportions) or raw counts. We denote by *logX_i_*(*t_ij_*) the logarithm transformation of microbiome abundances after zero imputation. The log-abundance trajectory of component A for individual *i* is denoted by *logX_iA_* = (*logX_iA_*(*t*_*i*1_, *logX*_*i*2*A*_,…, *logX_iA_*(*t_iL_i__*)) and the log-ratio trajectory between components A and B for individual *i* is given by:

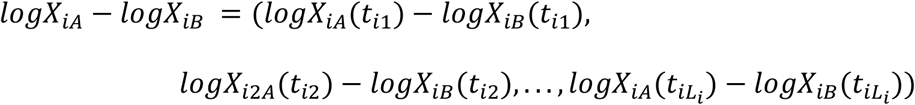

We summarize the log-ratio trajectories within two time points *l*_1_ and *l*_2_ as the integral of the log-ratio trajectory:

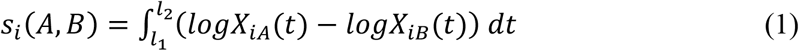

where the values of the log-ratio for *t* ∉ (*t*_*i*1_, *t*_*i*2_,…, *t_iL_i__*) are linearly interpolated.

We do not take the absolute value in equation (1) because the sign of the integral is informative: Positive values of *s_i_*(*A*, *B*) correspond to trajectories of component A above trajectories of component B, that is, larger relative abundances of A with respect to B, while negative values represent the opposite. Values of *s_i_*(*A*, *B*) around zero can represent similar abundances between A and B over time or a non-homogeneous trend between A and B within the observed region.

Another advantage of the summary *S_i_*(*A*, *B*) is computational. Since the integral is linear, *S_i_*(*A*, *B*) is equal to the difference between the integrals of log-transformed microbiome abundances of taxa A and taxa B:

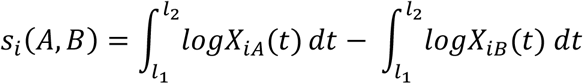

Thus, the number of integrals to be calculated is of the order of *K*, the number of taxa, instead of *K*(*K* – 1)/2, the number of pairwise log-ratios.

The log-ratio summary for the *n* subjects, *s*(*A*, *B*) = (*s*_1_(*A*, *B*),…, *s_n_*(*A*, *B*)), can be tested for association with the phenotype *Y* = (*Y*_1_,…, *Y_n_*) with a generalized linear model (glm) adjusted for some covariates Z:

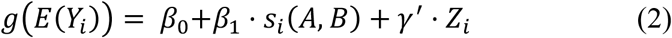

where *β*_0_ is the intercept, *β*_1_ is the regression coefficient for the log-ratio summary between components *A* and *B*, *Z* = (*Z*_1_, *Z*_2_,…, *Z_r_*) are non-compositional covariates and γ is the vector of regression coefficients for Z.

### Microbiome signature based on log-ratio analysis

To identify those log-ratios that are most associated with the outcome *Y*, we implement glm penalized regression on the log-ratio summaries for all pairs of taxa:

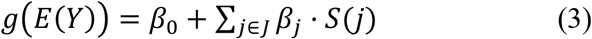

where *J* = {1,…,*K*(*K* – 1)/2} and *S*(*j*) = *s*(*j*_1_, *j*_2_) is the log-ratio summary of components *j*_1_ and *j*_2_ with (*j*_1_, *j*_2_) ∈ *J*_12_, the set of all possible combinations of pairs of components.

The regression coefficients in equation (3) are estimated to minimize a loss function *L*(*β*) subject to a penalization on the regression coefficients, *P*(*β*):

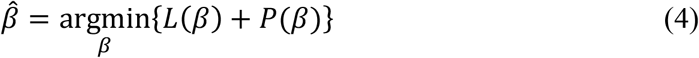

For the penalty term we consider the elastic-net, which combines the L1 and L2 norms: 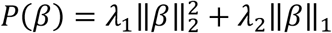. A common reparameterization of *P*(*β*) is *λ*_1_ = *λ*(1 – *α*)/2 and *λ*_2_ = *λα* where *λ* controls the amount of penalization and *α* the mixing between the two norms.

For the linear regression model the loss function is given by the residual sum of squares

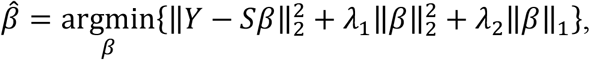

where *S* is the matrix of all log-ratio summaries and has dimension *n* by *K*(*K* – 1)/2. The expression of the optimization problem (4) for other models, like the logistic regression and the multinomial regression models, can be found in Friedman et al. (2010). Non-compositional covariates *Z* are previously modeled with Y and the fitted values are considered as “offset” in the penalized regression.

The result of the penalized optimization provides a set of selected pairs of taxa, those with a non-null estimated coefficient. For each individual, *i* ∈ {1,…, *n*}, the linear predictor of the generalized linear model (3), 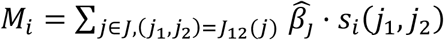, is the microbiome signature which is associated with the phenotype *Y_i_*. Because of the linearity of the integrals used as summaries of the log-ratio trajectories, the microbiome signature *M* can be rewritten in terms of the selected single taxa which is more interpretable than the selected pairs of components:

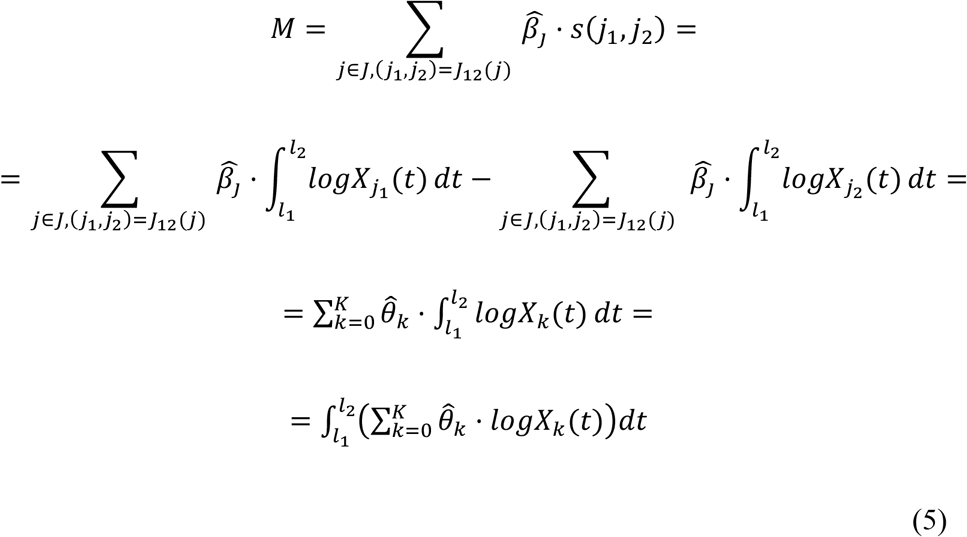

where 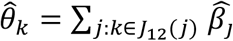, that is, the sum of the coefficients 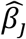 corresponding to a log-ratio that involves component *k*.

It can be proved that 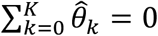 and thus, the microbiome signature *M* is the integral of the trajectory of a log-contrast function involving the selected taxa (those with 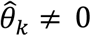). This ensures the invariance principle required for proper compositional data analysis and it facilitates the interpretation of the microbiome signature: Expression 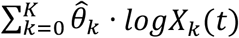 in (5) can be interpreted as a weighted balance between two groups of taxa, *G*_1_ and *G*_2_, the taxa with a positive coefficient vs those with a negative coefficient (Susin et al. 2020).

The package “coda4microbiome” (Calle and Susin, 2022) contains several functions that implement the proposed algorithms. The two main functions are `explore_lr_longitudinal()`, that implements the simple generalized linear model (equation 2), and `coda_glmnet_longitudinal()`, that performs penalized regression for the multivariable generalized linear model (equation 4). Additional functions are available like function `plot_signature_curves()` that provides a plot of the signature trajectories or `filter_longitudinal()` that filters those individuals and taxa with enough longitudinal information.

To illustrate the proposed approach and the R implementation we use data from the early childhood and the microbiome (ECAM) study (Bokulich et al. 2016). Metadata and microbiome data were downloaded from https://github.com/caporaso-lab/longitudinal-notebooks. Microbiome data, corresponding to 16S rRNA gene microbiota compositions sampled at regular intervals, were available in QIIME 2 qza file format (file ecam-table-genus.qza) and were transformed to R objects with function read_qza() of the R library qiime2R: https://github.com/jbisanz/qiime2R. Metadata (file ecam-sample-metadata.tsv) were in long format: multiple rows for individual, one for each time-point observation. Initially the data contained information on 43 child and 445 taxa at the genus level. We filtered those individuals and taxa with enough information for time-course profiling: we removed individuals with only one time-point observation and those taxa with less than 30 children (70% of individuals) with at least 3 non-zero observations over the follow-up period. After filtering, the data reduced to 42 children and 37 taxa.

## 3. Results

We demonstrate the proposed methodology with data from the “Early childhood and the microbiome (ECAM) study” that followed a cohort of 43 U.S. infants during the first 2 years of life for the study of their microbial development and its association with early-life antibiotic exposures, cesarean section, and formula feeding (Bokulich et al. 2016). Microbiome data were available for 43 child and 445 taxa at the genus level (Bokulich et al. 2018). After filtering those individuals and taxa with enough information for time-course profiling, the data were reduced to 42 child and 37 taxa. We focus on the effects of the diet on the early modulation of the microbiome by comparing microbiome profiles between children with breastmilk diet (bd) vs. formula milk diet (fd) in their first 3 months of life.

### Most important taxa

By implementing the pairwise log-ratio approach for longitudinal data (function `explore_lr_ongitudinal()`), we identified which taxa have more different dynamics between bd and fd children in the first three months of life. Table 1 provides the top 15 taxa with more discriminative dynamics between both diets.

**Table 1.** Taxa with most different abundances between the two diets groups during the first three months of life.

| Taxonomic assignment | More abundant group |
| --- | --- |
| "p_Proteobacteria;c_Gammaproteobacteria;o_Pasteurellales;f_Pasteurellaceae;g_Haemophilus" | bd |
| "p_Firmicutes;c_Bacilli;o_Bacillales;f_Staphylococcaceae;g_Staphylococcus" | bd |
| "p_Firmicutes;c_Clostridia;o_Clostridiales;f_Veillonellaceae;g_Veillonella" | fd |
| "p_Firmicutes;c_Clostridia;o_Clostridiales;f_Lachnospiraceae;g_Blautia" | fd |
| "p_Firmicutes;c_Clostridia;o_Clostridiales;f_Ruminococcaceae;g_1" | fd |
| "p_Firmicutes;c_Clostridia;o_Clostridiales" | fd |
| "p_Firmicutes;c_Clostridia;o_Clostridiales;f_Clostridiaceae" | fd |
| "p_Actinobacteria;c_Actinobacteria;o_Bifidobacteriales;f_Bifidobacteriaceae;g_Bifidobacterium" | bd |
| "p_Firmicutes;c_Clostridia;o_Clostridiales;f_Lachnospiraceae" | fd |
| "p_Bacteroidetes;c_Bacteroidia;o_Bacteroidales;f_Bacteroidaceae;g_Bacteroides" | bd |
| "p_Firmicutes;c_Bacilli;o_Lactobacillales;f_Lactobacillaceae;g_Lactobacillus" | bd |
| "p_Firmicutes;c_Erysipelotrichi;o_Erysipelotrichales;f_Erysipelotrichaceae;g_[Eubacterium]" | fd |
| "p_Firmicutes;c_Clostridia;o_Clostridiales;f_Lachnospiraceae;g_Coproccoccus" | fd |
| "p_Firmicutes;c_Clostridia;o_Clostridiales;f_Lachnospiraceae;g_Dorea" | fd |
| "p_Firmicutes;c_Bacilli;o_Lactobacillales;f_Enterococcaceae;g_Enterococcus" | fd |

**Table 2.**
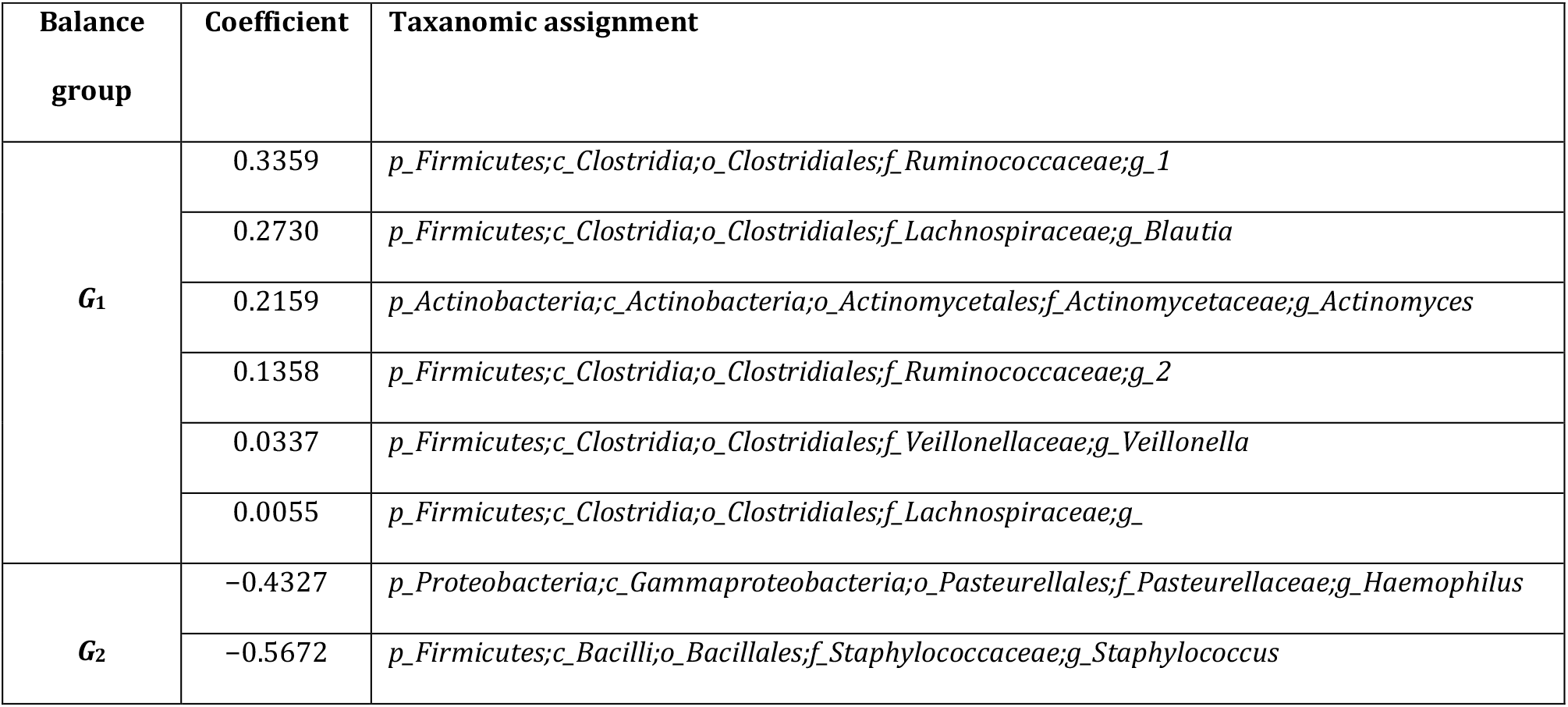
Taxa included in the microbiome signature that best discriminates between the two diet groups

| Balance group | Coefficient | Taxonomic assignment |
| --- | --- | --- |
| $G_1$ | 0.3359 | *p_Firmicutes;c_Clostridia;o_Clostridiales;f_Ruminococcaceae;g_1* |
|  | 0.2730 | *p_Firmicutes;c_Clostridia;o_Clostridiales;f_Lachnospiraceae;g_Blautia* |
|  | 0.2159 | *p_Actinobacteria;c_Actinobacteria;o_Actinomycetales;f_Actinomycetaceae;g_Actinomyces* |
|  | 0.1358 | *p_Firmicutes;c_Clostridia;o_Clostridiales;f_Ruminococcaceae;g_2* |
|  | 0.0337 | *p_Firmicutes;c_Clostridia;o_Clostridiales;f_Veillonellaceae;g_Veillonella* |
|  | 0.0055 | *p_Firmicutes;c_Clostridia;o_Clostridiales;f_Lachnospiraceae;g_* |
| $G_2$ | -0.4327 | *p_Proteobacteria;c_Gammaproteobacteria;o_Pasteurellales;f_Pasteurellaceae;g_Haemophilus* |
|  | -0.5672 | *p_Firmicutes;c_Bacilli;o_Bacillales;f_Staphylococcaceae;g_Staphylococcus* |

### Microbiome signature

The application of the proposed algorithm (with function `coda_glmnet_longitudinal()`) identified a microbiome signature with maximum discrimination accuracy between the two diet groups. The signature is defined by the relative abundances of two groups of taxa, *G*_1_ and *G*_2_, where *G*_1_ is composed of 6 taxa (those with a positive coefficient in the regression model) and *G*_2_ is composed of 2 taxa (those with a negative coefficient) (Table 1 and Figure 1). Group *G*_1_ is mainly dominated by three taxa within the order *Clostridiales* (family *Ruminococcaceae* (2) and gender *Blautia*) and one taxon within the gender *Actinomyces*. Two taxa (*g_Veillonella* and *f_Lachnospiraceae*) have a coefficient close to zero and will have a very small contribution to the signature. Group *G*_2_ is composed by two taxa within the genders *Haemophilus* and *Staphylococcus*. Note that the selected taxa within the microbial signature are among most important taxa according to the results of the pairwise log-ratio analysis (Table 1).

**Fig 1.**
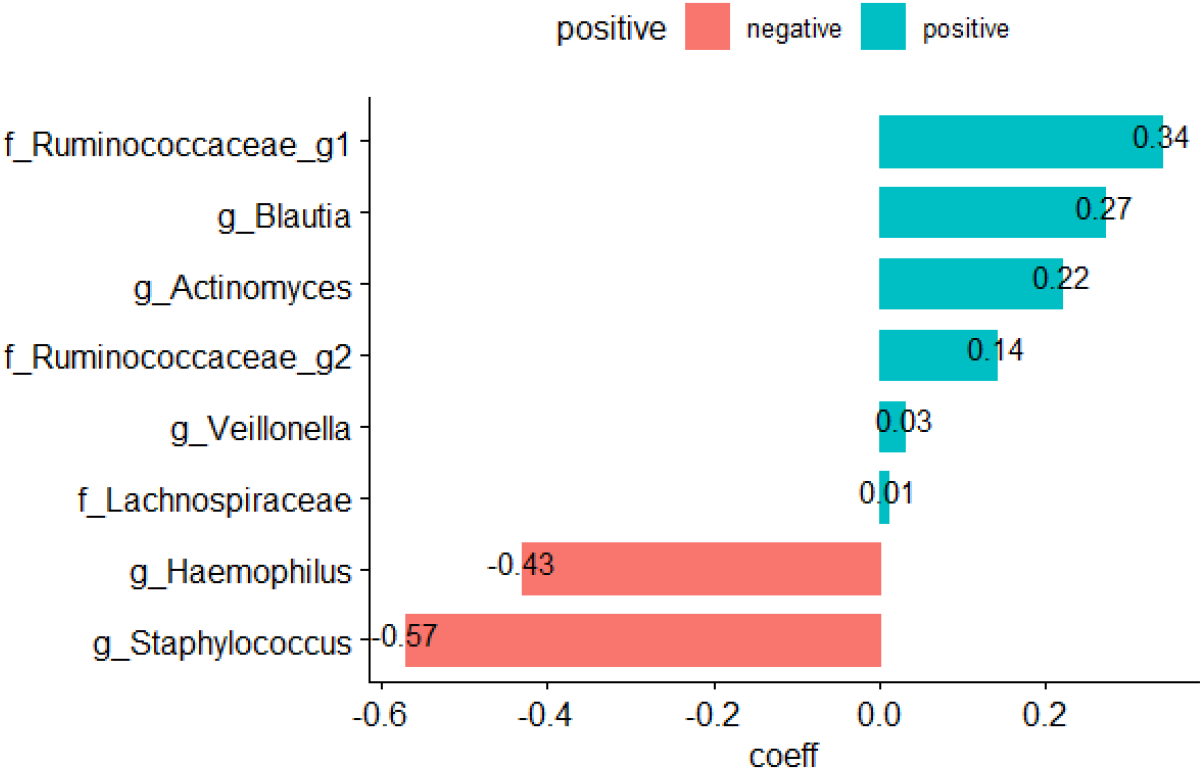
Taxa composing the microbiome signature that best discriminates between the two diet groups (green: positive coefficient and red: negative coefficient)

The trajectories of the microbial signature over the observed period are represented in Figure 2, where the color of the curves corresponds to the diet group. Each trajectory represents the relative mean abundances between the two taxa groups for each child. We can see that the two groups are clearly separated. Those children under breastmilk diet (in orange) usually have trajectories below zero, which means they have more relative mean abundance of *g_Haemophilus* and *g_Staphylococcus* with respect to the relative abundance of taxa in group *G*_1_, while children with formula milk diet (in blue) have more relative abundance of taxa in group *G*_1_ relative to *G*_2_.

**Fig 2.**
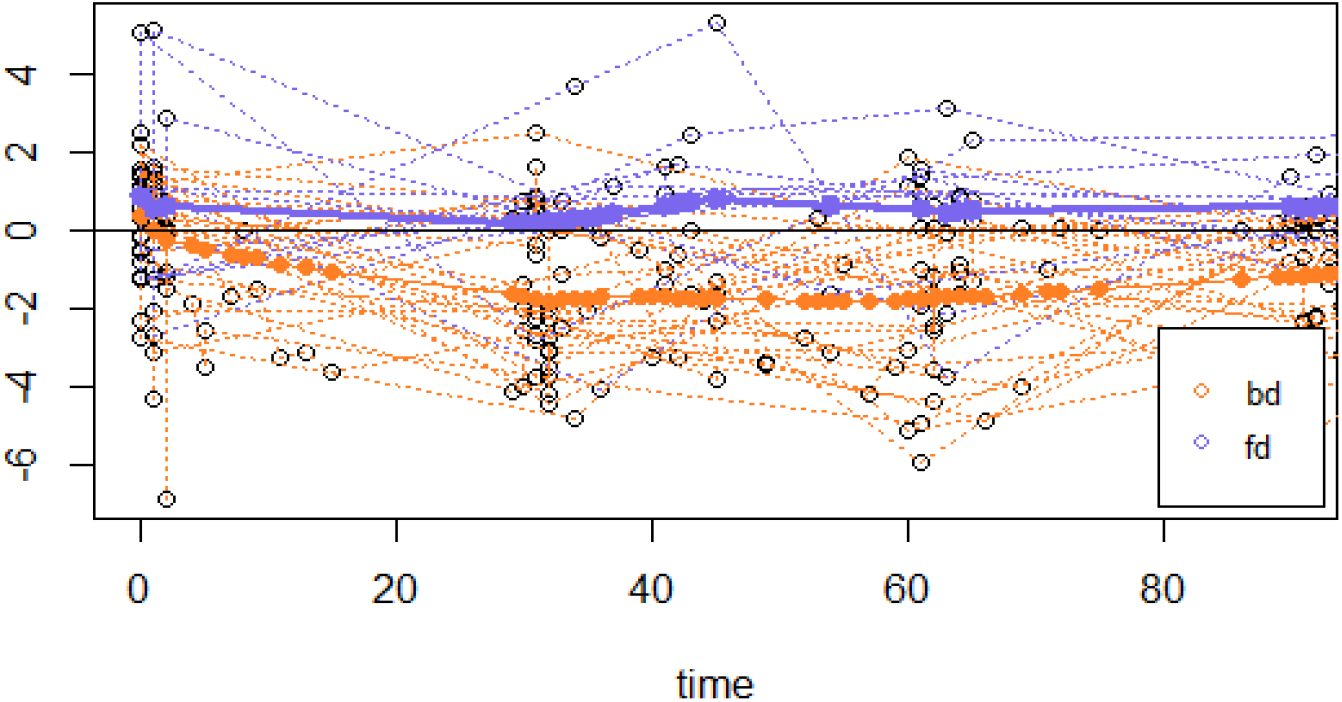
Relative abundance between group *G*_1_ and *G*_2_ during the first three months of life. Highlighted curves represent the mean value of the signature for each diet group (orange: breast milk diet, blue: formula milk diet)

Figure 3 displays the distribution of the microbial signature scores for the two diet groups and offers a visual assessment of the (apparent) discrimination accuracy of the signature. Quantitatively, the apparent discrimination accuracy of the signature (i. e. the AUC of the signature applied to the same data that was used to generate the model) is 0.96 and the mean cross-validation AUC is 0.74 (sd=0.10).

**Fig 3.**
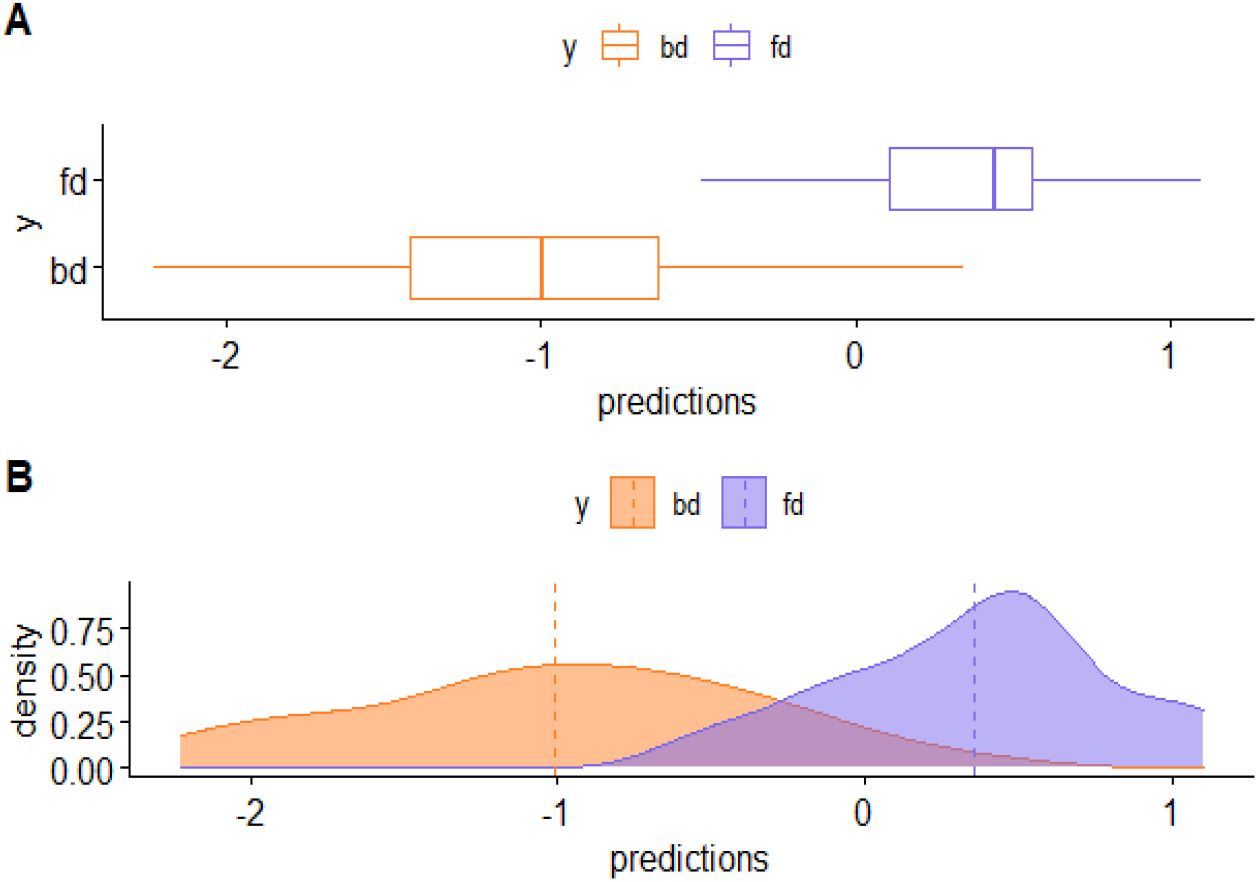
Distribution of the microbial signature scores for the two diet groups (orange: breast milk diet, blue: formula milk diet)

Both results, the pairwise analysis and the taxa selected in the microbial signature, are consistent with previous studies on the association of the infant gut microbiome composition and breastmilk feeding practices. In Fehr et al. (2020), *Haemophilus parainfluenzae* and *Staphylococcus* were found to be enriched with exclusive breastmilk feeding together with lower prevalence of *Actinomyces* at 3 months. *Lachnospiraceae* (*Blautia*) was enriched among infants who were no longer fed breastmilk. Similar results are reported in Laursen et al. (2016) where the duration of exclusive breastfeeding was negatively correlated with genera within *Lachnospiraceae* (e.g., *Blautia*) and genera within *Ruminococcaceae*. Positive correlations with exclusive breastfeeding were observed for *g_Bifidobacterium* and *Pasteurellaceae* (*Haemophilus*).

## 4. Discussion

Longitudinal microbiome studies, especially those focused on the human microbiome, have usually low resolution: the number of individuals is small, each individual has few observation times, the observations of the different individuals are not made at exactly the same time, the data are very variable, the expected behavior of the abundance trajectories is not linear or quadratic, etc. This makes it difficult to justify and implement a parametric modeling of trajectories and limits the use of models for longitudinal data (time series, mixed models). In this context, a description of the trajectories such as the one we propose, although less precise, allows to extract valuable information from the data as we have shown in the example. Other longitudinal data modeling strategies (Gerberg et al. 2012, Park et al. 2020, Silverman et al. 2018, Äijö et al. 2017) could be used in longitudinal microbiome studies with higher resolution such as laboratory or animal experimental studies.

The applicability of CoDA methods in microbiome studies has been limited by the difficulty in interpreting the obtained results. We hope that this work and the R package “coda4microbiome” will help to increase the use of these methods in this field.

## Acknowledgements

This work was partially supported by the Spanish Ministry of Economy, Industry and Competitiveness, Reference PID2019-104830RB-I00.

## Data Accessibility

The filtered data from the ECAM study is available as a data object in the “coda4microbiome” package.

